# Making Biorisk Measurable: A Bayesian Framework for Laboratory Risk Management

**DOI:** 10.64898/2026.03.09.710076

**Authors:** Dimiter Prodanov

## Abstract

Biosafety risk assessment traditionally relies on categorical scales embodied by the four WHO Risk Groups and biocontainment levels. Mapping such categories to quantitative metrics is an open problem for the field: the classifications are too coarse for operational decision-making, yet strictly probabilistic language remains inaccessible to most safety professionals, laboratory managers, and decision-makers. To bridge these gaps, the present work develops a quantitative Bayesian framework for laboratory risk management that combines WHO Risk Group classification as a prior with a Markov chain model of the incident-disaster escalation chain. Risk is reported on a log-risk scale that transforms multiplicative probabilities into additive quantities, mirroring the decibel scale in acoustics. The framework accommodates longitudinal updating with local incident data and quantifies the separate contributions of training, preventive maintenance, and inspection to system-level safety. Optionally, resource allocation recommendations providing auditable, evidence-based prioritisation can be issued. The framework is illustrated on synthetic BSL-3 scenarios and shifts the perspective of biorisk governance from static compliance assessment to dynamic risk and resource management.

## 1. Introduction

Biosafety risk assessment traditionally uses risk and control banding methodology and relies on categorical scales to classify hazard and risk [4]. Mapping such scales to quantitative metrics is an open problem for the field.

When viewing risk traditionally, frequentist approaches rely on statistical assemblies of outcomes that allows one to estimate underlying probabilistic structures. This approach is useful when the tracked outcomes are frequent and the consequences – not very adverse. On the other hand, high-impact adverse events, (e.g. “black swans”) can come from heavy-tailed probability distributions. These present themselves as rare events, where the sample sizes are very small but the consequences can be severe. Incidents and accidents, such as laboratory-acquired infections are rare events, making frequentist probability estimates based on historical data effectively impossible. On the other hand, Bayesian approach provides a way of objectifying and quantifying assumptions, which can be applied even in small sample size circumstances. Another advantage is the adaptation to the incoming data or evidence, which allows for refining the outcome probability distribution.

Quantifying risk also faces a fundamental communication challenge. Naive use of strictly statistical/probabilistic language will be incomprehensible for most safety professionals, laboratory managers and decision-makers who are not necessarily mathematically trained.

To bridge so-identified gaps the present work presents a quantitative framework that combines WHO Risk Group classification as a Bayesian prior on the risk with Markov chain representing system evolution. The problems framed above are not unique for biosafety. Therefore, addressing them systematically can have positive impact also on other fields and vice versa adapting approaches from other fields to the unique specifics of biosafety will have positive impact on biorisk management.

### 1.1. Evolution in WHO Laboratory Biosafety Guidelines

The transition from the third to the fourth edition of the WHO Laboratory Biosafety Manual represents a paradigm shift that directly motivates the quantitative framework presented in this work. Understanding this evolution is essential for positioning the methodology within current biosafety professional guidance.

#### WHO LBM3: The Prescriptive Era (2004)

The third edition of the WHO Laboratory Biosafety Manual, published in 2004, established a prescriptive approach that became a de facto global standard [4]. Laboratories were classified into four biosafety levels (BSL-1 to BSL-4), each with corresponding facility requirements, equipment specifications, and operational practices. This framework implicitly equated an agent’s Risk Group with the required Biosafety Level–a RG-3 agent necessitated a BSL-3 laboratory. While this approach provided clear, practical guidance, it had significant limitations. The manual covered basic laboratories (BSL-1/2), containment laboratories (BSL-3), and maximum containment laboratories (BSL-4) in distinct sections, each with prescribed codes of practice, facility designs, and operational requirements. This rigidity failed to account for variations in handled agents, procedures, personnel competence, or lab culture. As noted in the scholarly record, this led to the “misconception that the risk group of a biological agent directly corresponds to the biosafety level of a laboratory”. With time and practice some necessary mismatch between biocontainment level and the agent risk group became accepted – for example, RG 3 blood-borne viruses could be handled at containment level 2; the advent of unofficial BSL-2 sub-levels, etc.

#### WHO LBM4: The Risk-Based Paradigm (2020)

The fourth edition of the WHO Laboratory Biosafety Manual (LBM4), published in December 2020 after five years of development, fundamentally reconceptualized biosafety. Notably, LBM4 explicitly abolishes the direct correspondence between Risk Group and Biosafety Level (BSL), instead promoting an evidence-based, risk-based approach. LBM4 is structured as a core document accompanied by seven subject-specific monographs addressing risk assessment, laboratory design, biosafety cabinets, personal protective equipment, decontamination and waste management, biosafety program management, and outbreak preparedness. This modular structure allows facilities to implement “feasible and most effective combination of risk control measures based on their resources, experience, and local context”. Furthermore, the WHO LBM4 advocates for risk-based, case-by-case assessment rather than rigid classification [5].

## 2. Problem Formulation

Adoption of WHO LBM4’s risk-based approach as an authoritative guidance creates an urgent need for quantitative tools. The manual emphasizes that “a thorough, evidence-based and transparent assessment of the risks allows safety measures to be balanced with the actual risk”. However, it provides limited guidance on how to quantify risk, measure control effectiveness, or optimize resource allocation across competing interventions. This article presents a methodology that addresses three interconnected lab management problems:

1. **The rare event problem**: How to quantify risks when no local incidents have yet occurred.
2. **The communication problem**: How to make quantitative risks intuitive for decision-makers and lab workers.
3. **The optimization problem**: How to allocate limited safety budgets for maximum risk reduction.

### The calibration problem

The epidemiological burden of laboratory-acquired infections (LAIs) has been documented since Pike’s landmark survey of 3,921 cases across 159 agents [10], yet quantitative per-procedure risk estimates remain elusive due to a persistent denominator problem: no survey has provided the number of procedures performed as a base for incidence calculation [12, 16]. Historically, infection rates in UK clinical laboratories fell from 82.7 to 16.2 cases per 100,000 person-years between 1988-89 and 1994-95 – a five-fold reduction attributable to systematic adoption of biosafety interventions – yet no mathematical framework exists to decompose this improvement into training, maintenance, and inspection contributions [13, 14]. BSL-3 specific surveillance found different contributions, such as that 73% of LAIs in containment laboratories arose during routine microbiology activities rather than exceptional events, consistent with our model’s anchoring of baseline risk to the *S*_0_ → *S*_1_ procedural deviation pathway rather than to catastrophic equipment failure [12]. Baron and Miller (2008) reported 3.5 LAIs per 1,000 full-time employees per year in hospital diagnostic laboratories, corresponding to a per-procedure log-risk of *L* ≈ 6.5 under a standard assumption of 10,000 procedures per employee per year – a value our model reproduces at baseline without parameter fitting [11].

The structural deficit in LAI surveillance infrastructure has been repeatedly acknowledged: as of 2016, no national or international registry existed to systematically record LAI exposures, and the level of detail in published case reports varies considerably [15]. ABSA International subsequently developed a searchable database of peer-reviewed LAI case reports [15], yet even this resource captures only reported cases with no procedural denominator, leaving per-procedure risk unquantified.

Accidental pathogen release is also a rare event. On the release side, Klotz and Sylvester estimated an annual accidental release probability of 0.003 per high-biosafety lab (2003-2017), yielding a cumulative five-year community release risk of 15.8% and pandemic risk of approximately 2.5% [29].

Given the above background, the objective of the present paper focused on two interconnected aspects: on one hand it aims to exemplify utility of quantitative approaches; on the other hand the paper also aims to start the debate on acceptable parameter choices.

## 3. Mathematical Framework

The framework has four components, described intuitively here and specified mathematically in Appendix A.

### (1) The log-risk scale

We report laboratory safety as a single number *L*, defined so that higher values mean safer conditions – exactly as decibels measure sound, where higher dB means louder. A value of *L* = 7 means one laboratory accident is expected every 10^7^ procedures; *L* = 6 means ten times more risk. The scale maps directly onto WHO Risk Groups: RG-1 (*L* ≈ 2), RG-2 (*L* ≈ 4), RG-3 (*L* ≈ 6), RG-4 (*L* ≈ 8). We do not make a distinction between pathogens and genetically modified organisms (GMOs). Doubling the mean time between accidents adds approximately 0.3 to *L*, mirroring the 3 dB rule in acoustics. Two companion metrics, *L*_*M*_ and *L*_*I*_, isolate the maintenance and inspection channels respectively, enabling clean, non-overlapping return-on-investment calculations.

### (2) The escalation chain

Laboratory operations are modelled as a five-state system (Fig. 1): normal (*S*_0_), minor deviation (*S*_1_), serious equipment issue (*S*_2_), critical / containment threatened (*S*_3_), and disaster (*S*_4_, absorbing). Each procedure moves the system one step according to transition probabilities that depend on the current intervention levels. The three panels of Fig. 2 show which transitions each intervention controls.

**Figure 1:**
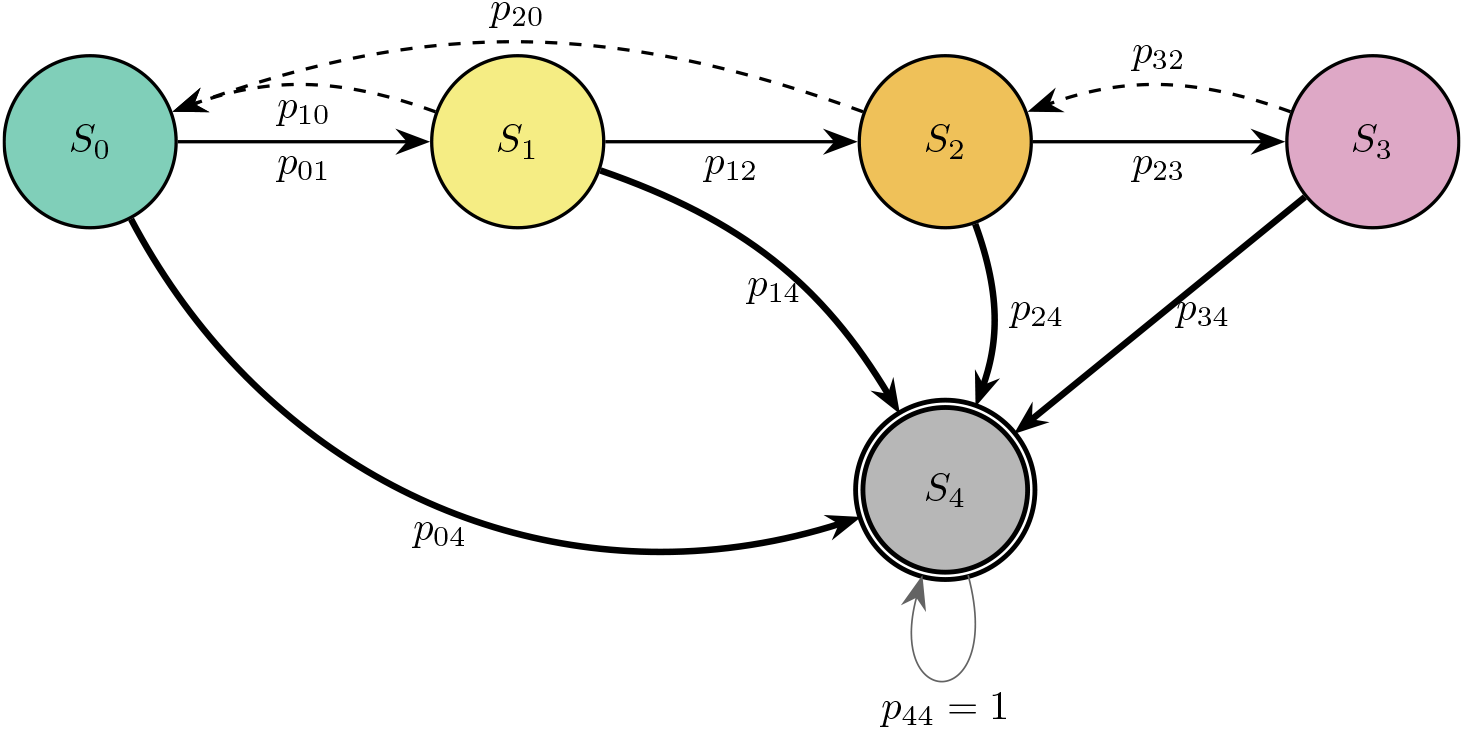
Markov Chain Escalation Diagram Legend: *S*_0_ : Normal operation (all systems functional) *S*_1_ : Minor issues (procedural deviations, fatigue) *S*_2_ : Serious issues (equipment malfunction) *S*_3_ : Critical (containment threatened/compromised) *S*_4_ : DISASTER (actual release/exposure)

**Figure 2:**
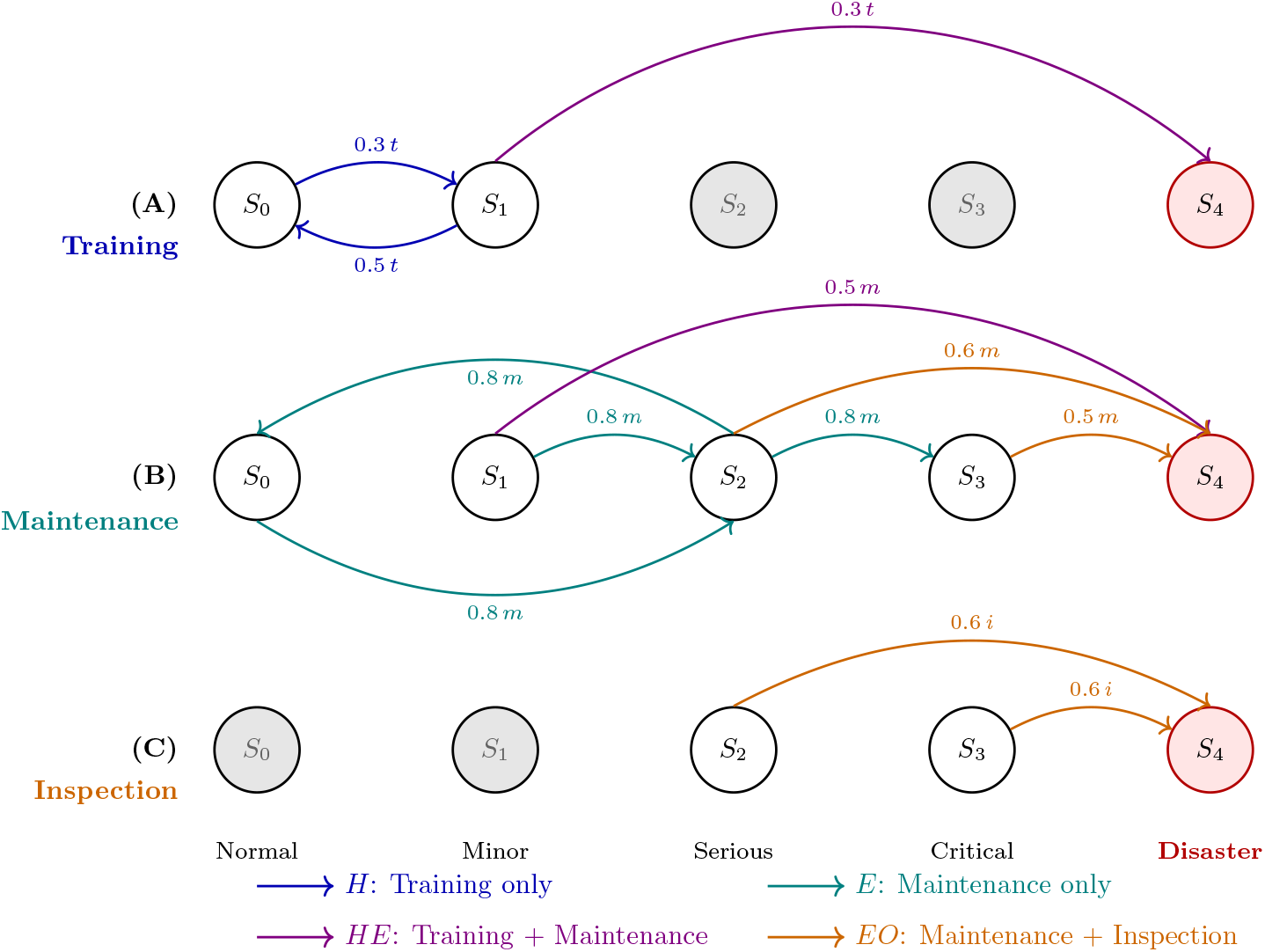
Markov chain paths: training, maintenance, inspection. Each panel highlights the edges modulated by one intervention; grey arrows are inactive for that channel. (A) Training acts on class *H* edges (*S*_0_ ↔ *S*_1_) and the *HE* shortcut (*S*_1_ → *S*_4_). (B) Maintenance acts on class *E* edges, including spontaneous equipment failure *S*_0_ → *S*_2_, and on the dual-class *HE*/*EO* edges. (C) Inspection acts exclusively on class *EO* edges (*S*_2_ → *S*_4_, *S*_3_ → *S*_4_). Edge labels give the weight coefficent (*t =* training, *m =* maintenance, *i* inspection effect).

### (3) Intervention effects

Training reduces the probability of a minor procedural deviation (*S*_0_ → *S*_1_), with diminishing returns above 50-60 hours (Hill function, Appendix A.4). Maintenance effectiveness depends on both scheduled frequency and the fraction of sessions actually completed; missed sessions create compounding vulnerability windows captured by an exponential gap penalty (Appendix A.5). Inspection produces a threshold effect: facilities scoring below 70/100 receive no risk reduction; those crossing the threshold gain Δ*L*_*I*_ = 0.600 logs on the inspection channel (Appendix A.6).

The edge *p*_04_ represents a Natech (natural hazard-triggered) disaster path (e.g. fire, flood, earthquake) and is parameterised separately from internal biosafety failures.

### (4) Bayesian updating

When local records are available, the framework updates its parameter estimates using Markov Chain Monte Carlo (MCMC). Three observable event types drive the update: near-misses, light incidents, and disasters. Crucially, the three composite risk metrics (*L, L*_*M*_, *L*_*I*_) are recoverable even when individual parameters (such as training hours) are not uniquely identifiable from the data (Section 4).

## 4. Results

To estimate the laboratory’s true training, maintenance, and inspection levels from the observed incident data, the model uses Markov Chain Monte Carlo (MCMC), an iterative estimation technique analogous to calibration by successive approximation: rather than solving directly for a single answer, the algorithm proposes a candidate parameter set, evaluates how well it explains the observed near-misses and incidents, and updates accordingly, repeating this cycle thousands of times until the proposals stabilise around the range of values best supported by the data. The output is not a single point estimate but a full uncertainty distribution for each parameter, analogous to a measurement uncertainty budget rather than a single calibrated value.

The model was evaluated across a range of intervention scenarios to assess its output and validate the underlying assumptions. Under the illustrative parameter set adopted in the paper, the model produced the following demonstrative outputs:

- Under the assumed cost and effect-size parameters, a laboratory investing $2,000-$4,000 in additional training (40-60 hours) would be projected to recover that cost in annual risk reduction within four months, corresponding to a modeled ROI of 2.6-2.9×.
- Within the model, maintaining equipment consistently (90,% schedule adherence) at modest frequency is projected to outperform highfrequency maintenance with poor compliance by a factor of ∼ 9 in effectiveness, at comparable or lower cost.
- Given the assumed inspection cost structure, a single compliance inspection that achieves a passing score (≥ 70*/*100) at a one-time cost of $30,000 is projected to produce the highest dimensionless return of any modeled intervention (0.020 Δ*L*_*I*_ per $1,000).
- Under the illustrative cost assumptions, a $100,000 annual safety budget, allocated optimally across all three interventions, is projected to reduce expected annual disaster cost from $58,867 to $24,232 – a 58.8,% reduction in this scenario.

These figures should be read as illustrations of the framework’s internal decision logic and utility under a specific parameterization; absolute magnitudes will shift under alternative input assumptions, whereas the qualitative patterns discussed below (adherence dominating frequency, diminishing training returns, inspection threshold effects) are the more robust contribution of this analysis.

### 4.1. Maintenance Effectiveness: Frequency vs. Adherence

The maintenance model was designed to test the hypothesis that schedule adherence dominates raw frequency. Table 1 shows the maintenance effect (in log-risk units) and resulting system log-risk for five representative scenarios, holding training constant at 50 hours and inspection at 75 (passing).

**Table 1:**
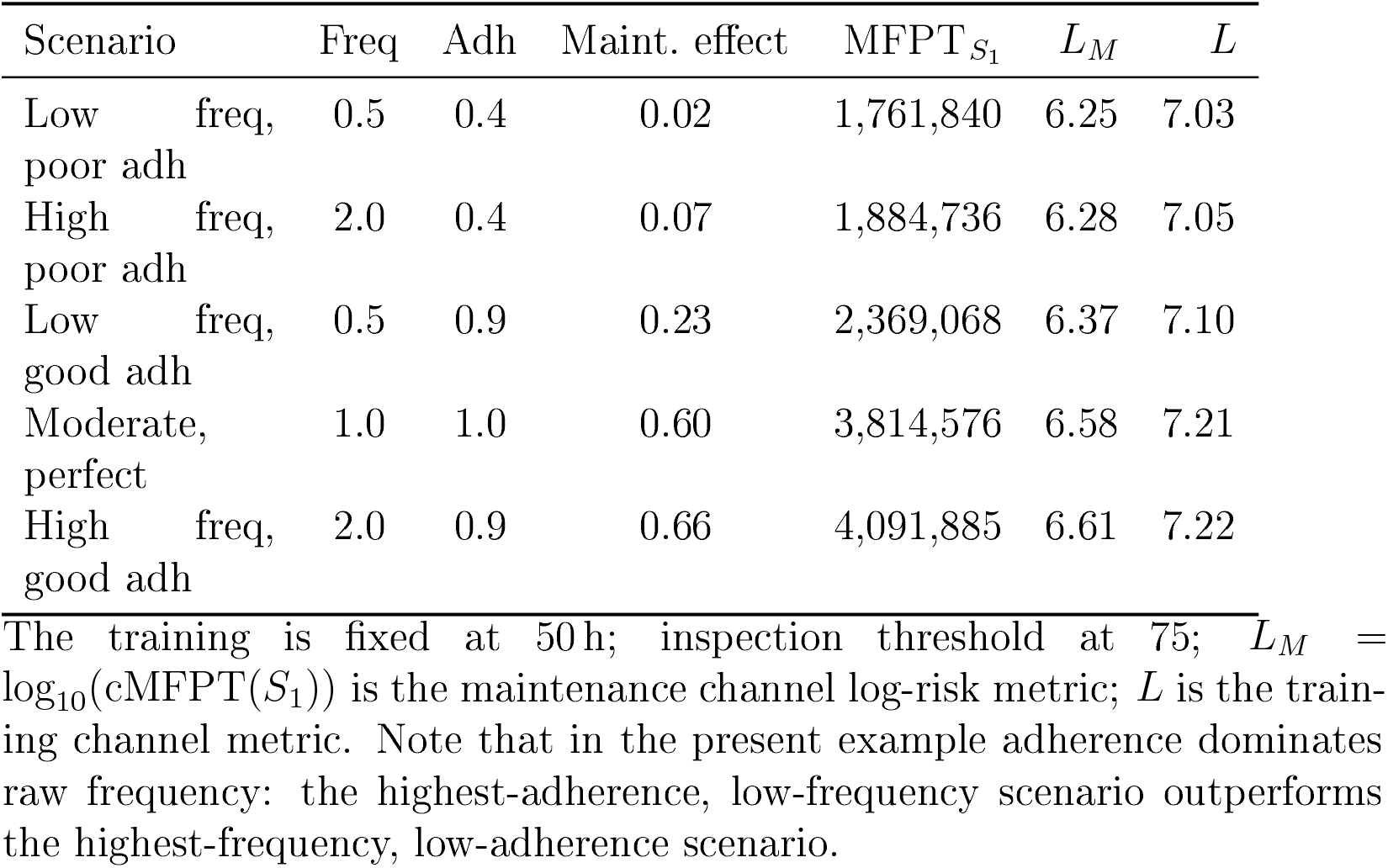
Maintenance effectiveness across different frequency and adherence scenarios.

| Scenario | Freq | Adh | Maint. effect | MFPT$_{S_1}$ | $L_M$ | $L$ |
| --- | --- | --- | --- | --- | --- | --- |
| Low freq, poor adh | 0.5 | 0.4 | 0.02 | 1,761,840 | 6.25 | 7.03 |
| High freq, poor adh | 2.0 | 0.4 | 0.07 | 1,884,736 | 6.28 | 7.05 |
| Low freq, good adh | 0.5 | 0.9 | 0.23 | 2,369,068 | 6.37 | 7.10 |
| Moderate, perfect | 1.0 | 1.0 | 0.60 | 3,814,576 | 6.58 | 7.21 |
| High freq, good adh | 2.0 | 0.9 | 0.66 | 4,091,885 | 6.61 | 7.22 |

The training is fixed at 50 h; inspection threshold at 75; $L_M = \log_{10}(\text{cMFPT}(S_1))$ is the maintenance channel log-risk metric; $L$ is the training channel metric. Note that in the present example adherence dominates raw frequency: the highest-adherence, low-frequency scenario outperforms the highest-frequency, low-adherence scenario.

Several patterns emerge from these results. First, **adherence dominates frequency**: a program with low frequency (0.5/month) but good adherence (90 %) achieves *L*_*M*_ = 6.37, while a program with high frequency (2.0/month) but poor adherence (40 %) achieves only *L*_*M*_ = 6.28 – a reversal confirming that consistency matters more than intensity.

Second, the **gap penalty** produces realistic non-linearities. Improving adherence from 40 % to 90 % at fixed frequency 2.0/month raises the maintenance effect from 0.07 to 0.66 logs – a factor of ∼9, far exceeding the proportional improvement in adherence, reflecting compounding vulnerability windows created by missed sessions.

Third, **diminishing returns to frequency** are evident at perfect adherence: doubling frequency from 0.5 to 1.0/month raises *L*_*M*_ by 0.21 (6.37 → 6.58), while a further doubling to 2.0/month adds only 0.03 (6.58 → 6.61). This saturation matches engineering experience.

The overall spread of *L*_*M*_ across scenarios is 0.37 logs (MFPT ratio 2.3 ×). The corresponding training channel metric *L* varies from 7.03 to 7.22 across the same scenarios, confirming that maintenance improvements propagate into system-level risk but are correctly attributed to the *L*_*M*_ axis rather than *L*.

### 4.2. Bayesian Updating with Simulated Data

To test the MCMC updating framework, a *moderate-safety BSL-3 analogue* scenario was specified, representative of a functional but not optimally resourced facility:

- Training: 35 hours
- Maintenance frequency: 0.8/month
- Maintenance adherence: 60 %
- Inspection score: 55 (below passing threshold)

The simulation uses a throughput of *λ* = 500 procedures per month; 36 months of operation therefore corresponds to 18,000 procedures total. MFPT-based metrics are reported in calendar months by dividing procedure-scale cMFPT values by *λ*. Three observable event types were recorded per month: near-misses (*S*_0_ → *S*_1_), light incidents (*S*_2_ → *S*_3_ escalations), and disasters (*S*_4_).

This configuration yields true metrics of *L* = 4.29, *L*_*M*_ = 3.37, and *L*_*I*_ = 2.25 (monthly scale). Over 36 months at *λ* = 500 procedures/month, the simulation produced 6 near-misses and 17 light incidents with 0 disasters – a realistic profile for a moderately resourced facility operating below optimal safety standards.

The MCMC chain was run for 20,000 iterations. The first 5,000 (the *burn-in*) were discarded, since early iterations are still anchored to the arbitrary starting values rather than to the data, much as the first readings from an instrument still equilibrating are excluded from a calibration record. Of the remainder, only every fifth iteration was retained (*thinning*), since consecutive iterations are highly autocorrelated and retaining all of them would overstate the effective sample size. The parameter ranges considered plausible before seeing any data (*weakly informative priors*, Section 4.2) were deliberately broad, ensuring the data – not the analyst’s starting assumptions – determine the final estimates whenever the data are sufficiently informative.

The *acceptance rate* of 0.731 (73.1 % of proposed updates were adopted) serves as a mixing diagnostic, analogous to a process-control check confirming the estimation procedure is sampling efficiently rather than stalling (acceptance rates persistently above ∼90 %) or producing unstable, poorly-explored estimates (rates below ∼10-20 %). A rate of 0.731 falls within the range conventionally regarded as indicating efficient exploration of the parameter space, supporting confidence that the reported uncertainty bounds – e.g. the 95 % credible interval for training hours, [35.4, 154.5] – reflect genuine data-driven uncertainty rather than a computational artefact. Posterior summaries are shown in Table 2.

**Table 2:** Simulated 36-month evolution of laboratory safety scores.

| Parameter | True | Mean | Std | 2.5 % | 97.5 % | CI covers? |
| --- | --- | --- | --- | --- | --- | --- |
| Training (hours) | 35.0 | 72.2 | 28.5 | 35.4 | 154.5 | NO |
| Maintenance frequency | 0.8 | 1.0 | 0.5 | 0.4 | 2.2 | YES |
| Maintenance adherence | 0.6 | 0.4 | 0.2 | 0.1 | 0.8 | YES |
| Inspection score | 55.0 | 43.7 | 30.9 | 0.0 | 98.0 | YES |
| $L$ (monthly) | 4.29 | 4.35 | 0.04 | 4.27 | 4.44 | YES |
| $L_M$ (months) | 3.37 | 3.51 | 0.12 | 3.33 | 3.78 | YES |
| $L_I$ (months) | 2.25 | 2.38 | 0.27 | 2.17 | 2.90 | YES |
| Acceptance rate: 0.731 (20,000 iterations, burn-in 5,000, thinning 5) |  |  |  |  |  |  |

Posterior summaries from MCMC updating with 36 months of simulated data (moderate-safety BSL-3 analogue; $\lambda = 500$ proc/month, 18,000 procedures total; 6 near-misses, 17 light incidents, 0 disasters observed). MFPT-based metrics are reported in calendar months ( $\text{cMFPT}_{\text{proc}}/\lambda$ ). Note that individual practices remain variable, but overall safety score remains reliable.

Several observations are noteworthy. First, **training is the hardest parameter to identify**: its posterior mean (72.2 h) is pulled strongly toward the prior mean (50 h) and the 95 % credible interval [35.4, 154.5] does not cover the true value of 35 h. This reflects a fundamental identifiability challenge: near-miss counts are driven by *P*_01_, but many combinations of training hours and base rates are consistent with the observed 6 near-misses over 18,000 procedures.

Second, **maintenance and inspection parameters are better identified** from the light-incident channel (*S*_2_ → *S*_3_): the 17 observed light incidents constrain the *S*_2_ → *S*_3_ transition rate and, through the weight structure, the maintenance effect. Both maintenance parameters and inspection have credible intervals covering the true values.

Third, and most importantly, **all three risk metrics are well recovered** despite parameter-level uncertainty. The posterior mean for *L* is 4.35 (true: 4.29), for *L*_*M*_ is 3.51 (true: 3.37), and for *L*_*I*_ is 2.38 (true: 2.25), all with narrow credible intervals relative to their parameter posteriors. This illustrates a key property of the framework: composite risk metrics are identifiable even when individual intervention parameters are not, because the likelihood surface is flat in parameter space but informative in metric space.

The inspection parameter shows a similarly wide posterior (Std = 30.9, 95% CI = [0.0, 98.0]) for the same underlying reason: with only 17 light incidents observed over 36 months, the likelihood surface constrains the composite metric *L*_*I*_ tightly (Std = 0.27) while leaving the underlying inspection score itself comparatively under-determined. This is a direct illustration of the framework’s core identifiability property – composite risk metrics are well recovered even when individual parameters, including inspection score, are not.

### 4.3. Cost-Effectiveness Analysis

Using the low-resource baseline (training = 20 h, freq = 0.5/month, adh = 50 %, insp = 50; *L* = 6.93, *L*_*M*_ = 5.93, *L*_*I*_ = 4.90, annual disaster cost $58,867) we conducted ROI analysis across the three intervention channels separately, using the three-metric decomposition.

#### Training and Maintenance ROI (Fig. 4)

Training yields the highest monetary return at 40-60 h ($5.7-10.5k/yr at ROI 2.9-2.6×), with saturation above 80 h. For maintenance, adherence dominates frequency: the highest adherence scenario (adh = 0.95, freq = 2.0) achieves 60 % better cost-efficiency than low-adherence alternatives.

#### Inspection

The inspection effect saturates at the 70-point threshold; all scores above 70 yield identical Δ*L*_*I*_ = 0.600 at identical cost ($30,000). At 0.0200 Δ*L*_*I*_/$1k, crossing the threshold offers the highest dimensionless return of any single intervention. The binary nature of the threshold formalises regulatory pass/fail inspection logic.

### 4.4. Optimal Budget Allocation

The optimal configuration within a $100,000 budget was: training = 120 h, freq = 2.0/month, adh = 95 %, insp = 70 (crossing threshold), at a total cost of $73,000. This generates an annual benefit of $34,159 (ROI = 0.5×) and reduces expected annual disaster cost from $58,867 to $24,232 – a **58.8 % reduction**. The budget is not fully exhausted because no additional combination within the remaining $27,000 improves total benefit, reflecting saturation of all three intervention functions at high investment levels.

### 4.5. Sensitivity to Incident Data

To assess how the model responds to actual incidents, a second simulation was run with the same typical scenario but introducing a single near-miss (state *S*_2_) in the 36-month period. The posterior for maintenance adherence shifted from 0.50 to 0.61, and the credible interval narrowed slightly. With one actual incident (state *S*_4_), the posterior for log-risk shifted downward (higher risk) and all parameters updated toward values consistent with the observed event.

This output confirms that the MCMC framework correctly updates objective beliefs in response to evidence, and that the priors are sufficiently weak to allow the data to dominate when informative.

1. **Maintenance:** Adherence dominates frequency. High adherence (90 %) at low frequency outperforms low adherence (40 %) at high frequency by Δ*L*_*M*_ = 0.09 logs. Optimal maintenance achieves a 2.3× improvement in 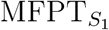 over worst-case maintenance.
2. **Training:** Best cost-efficiency at 40-60 h (ROI 2.6-2.9×, payback *<* 0.4 yr). Saturation above 80 h.
3. **Inspection:** Crossing the 70-point threshold yields Δ*L*_*I*_ = 0.600 (0.0200 Δ*L*_*I*_/$1k) – the highest dimensionless return per dollar of any intervention.
4. **Bayesian updating:** The three-observable likelihood recovers all three composite metrics (*L, L*_*M*_, *L*_*I*_) with credible intervals covering true values, even when training hours are individually non-identifiable from near-miss data alone.
5. **Optimal allocation:** A $100,000 budget achieves a 58.8 % reduction in expected annual disaster cost ($58,867 → $24,232).

These results validate the model’s behavior and support its use as a tool for biosafety risk assessment and resource allocation. The adherence-beats-frequency insight has immediate practical implications for laboratory management, suggesting that efforts to improve maintenance consistency may yield greater returns than simply increasing scheduled frequency.

## 5. Worked Example

Proposed framework matches a laboratory’s day-to-day safety practices to a single decibel safety score. Intuitively, a higher score always means a safer laboratory, while a lower score means more risk. The tool starts with the WHO Risk Group of the pathogen/ GMO being handled as the best available information and treats it as a sensible starting estimate of risk. From there, it adjusts that estimate using three items a laboratory manager already tracks in routine operations: (i) how much staff training has been completed, (ii) how consistently scheduled equipment maintenance has actually been carried out, and (iii) the facility’s most recent compliance inspection score. No specialized statistical knowledge is required to supply these inputs; they are typically present in the lab records already kept for safety compliance purposes.

Once these numbers are entered, the model produces three outputs: one overall safety score for the facility plus two additional scores that isolate the contribution of maintenance and inspection specifically. This separation matters because it tells a manager not just *how safe* the facility is overall, but *which lever to pull* to improve it fastest – for example, whether money is better spent on additional training, on tightening maintenance adherence, or on preparing for a compliance inspection. As real incidents or near-misses are recorded over time, hopefully in small numbers, the tool automatically refines its estimates, the way a weather forecast sharpens as new observations come in; a laboratory with zero recorded incidents is not assumed to be zero-risk, but instead retains a baseline evidence-based estimate drawn from its Risk Group and practices.

In practice, a manager can therefore be advised on an up-to-date safety readout for reporting, and a cost-based ranking of which improvement to implement first.

### 5.1. Applying the Framework in Practice

The five-step walk-through in this section applies this logic to a hypothetical BSL-3 laboratory. Consider a BSL-3 laboratory manager who wants to evaluate their current safety level. The process unfolds in five steps and matches closely the PDCA practice illustrated in Fig. 3.

**Figure 3:**
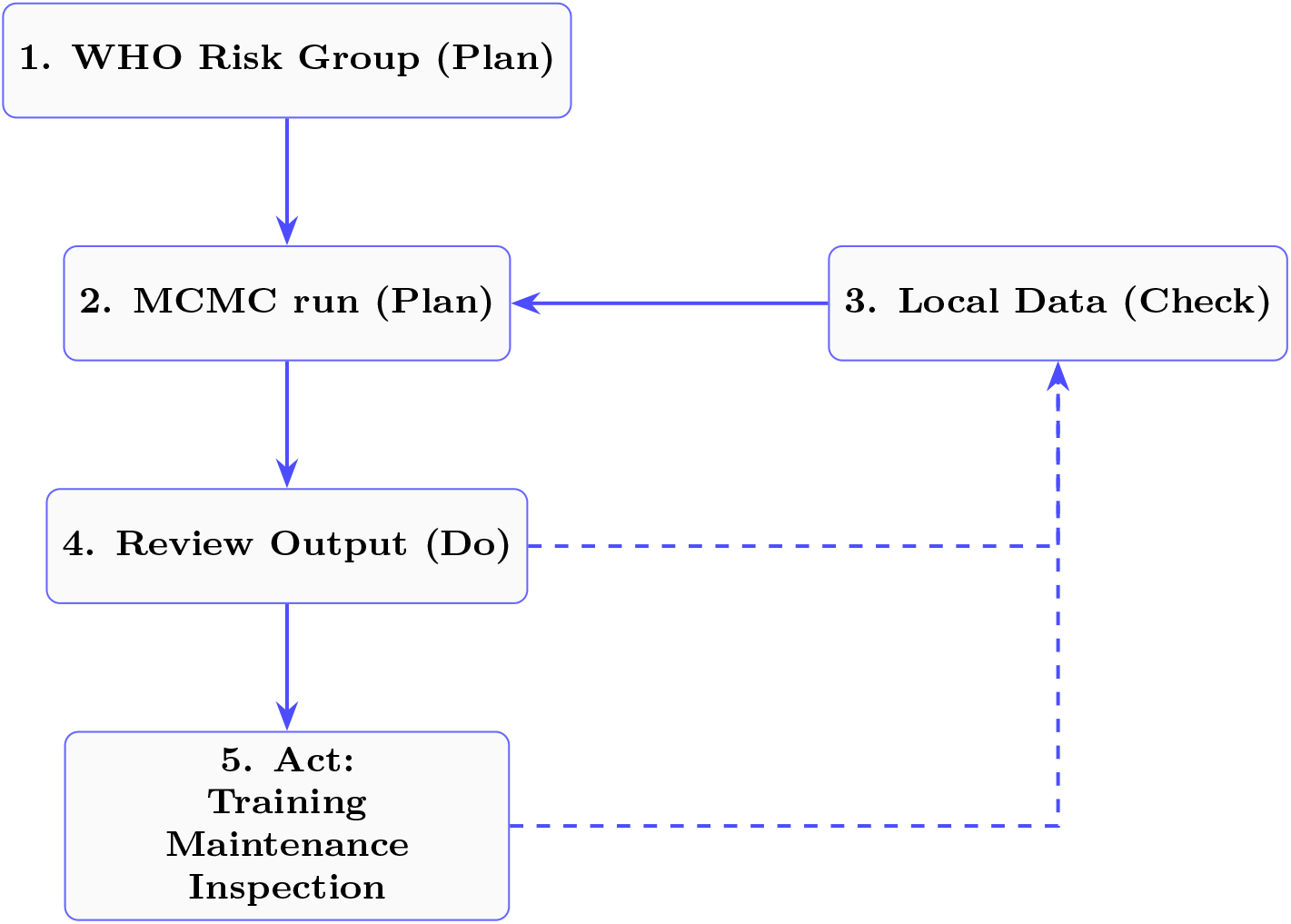
Laboratory manager workflow mapped to PDCA cycle. The framework implements Plan-Do-Check-Act through (1) Risk Group prior (Plan), (2) MCMC risk computation (Plan), (3) incident data collection (Check), (4) output review (Do), and (5) intervention execution (Act). Dashed feedback enables continuous improvement.

**Figure 4:**
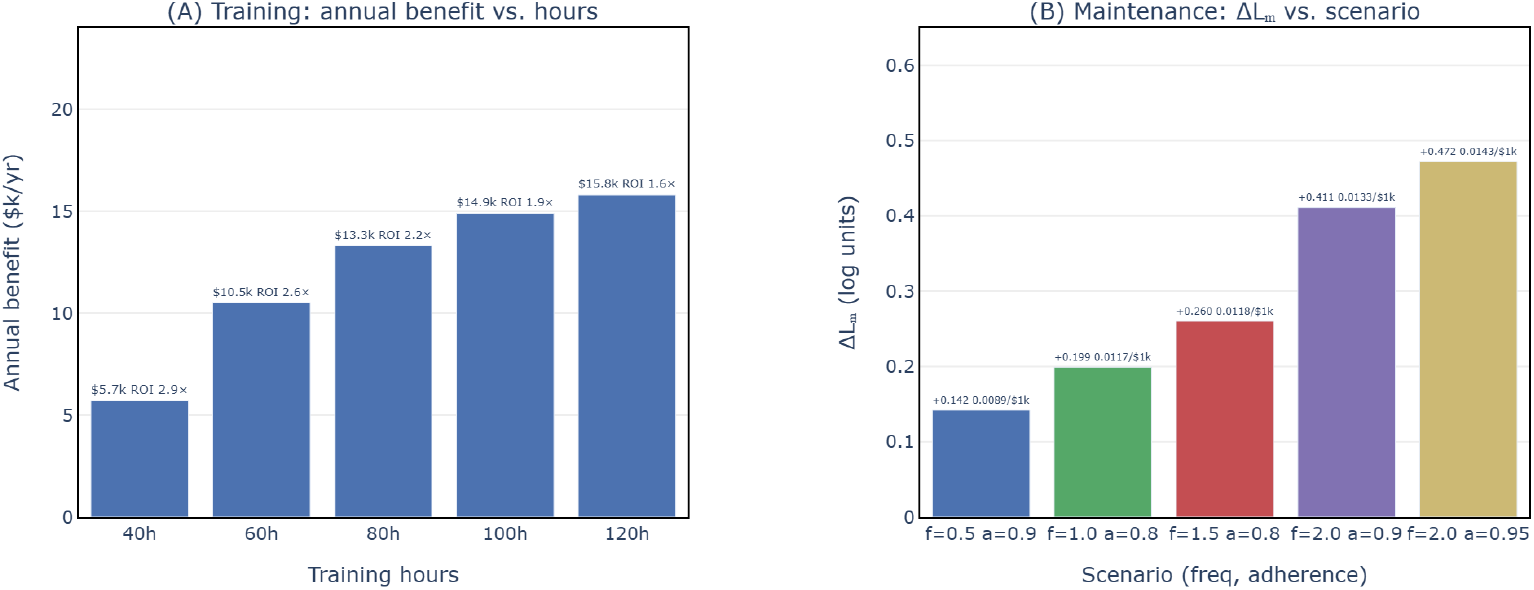
Cost-effectiveness of training (A) and maintenance (B) interventions from a low-resource baseline (20 h training, freq = 0.5/mo, adh = 50 %). Bar labels show absolute gain (annual benefit in $/yr for training; Δ*L*_*M*_ log-risk improvement for maintenance) and cost-efficiency per $1,000 invested. Colour coding in panel (B) distinguishes the five adherence-frequency combinations.

#### Step 1: Establish the starting point

The manager identifies the WHO Risk Group of the agents handled (e.g., RG-3), which sets an initial log-risk estimate of approximately *L* ≈ 6 before any facility-specific information is considered.

#### Step 2: Record current practices

The manager records three numbers already available from routine operations: annual training hours per staff member (e.g., 35 h), maintenance schedule adherence (e.g., 60 % of scheduled sessions completed), and the most recent inspection score (e.g., 55/100).

#### Step 3: Compute the composite risk score

These three inputs are entered into the model, which returns three numbers: the overall system log-risk *L*, and two channel-specific scores *L*_*M*_ (maintenance) and *L*_*I*_ (inspection). The interpretation is given in Table 3. In this example, the model returns *L* = 4.29, *L*_*M*_ = 3.37, *L*_*I*_ = 2.25, flagging inspection as the weakest link since the facility’s score of 55 falls below the 70-point compliance threshold.

**Table 3:** Operational interpretation of log-risk values.

| $L$ | Approx. meaning | Typical association |
| --- | --- | --- |
| 2 | 1 incident per 100 procedures | RG-1 agents, minimal containment |
| 4 | 1 incident per 10,000 procedures | RG-2 agents, standard BSL-2 practice |
| 6 | 1 incident per 1,000,000 procedures | RG-3 agents, well-run BSL-3 facility |
| 8 | 1 incident per 100,000,000 procedures | RG-4 agents, maximum containment |

#### Step 4: Collect incident data over time

Over the following months, the manager logs any near-misses (e.g., a procedural deviation caught before escalation) or incidents. Even a handful of such records – six near-misses and no incidents over three years, allow the model to refine its risk estimates via Bayesian updating, narrowing uncertainty without requiring a large historical dataset.

#### Step 5: Act on the ROI ranking

The model’s ROI analysis indicates that, for this facility, crossing the inspection threshold (raising the score from 55 to 70) offers the highest return per dollar invested ($30,000 for Δ*L*_*I*_ = 0.600), ahead of additional training or more frequent maintenance. The manager therefore prioritizes inspection readiness as the most cost-effective next investment.

## 6. Alignment with International Biorisk Management Standards

The proposed framework is designed as a quantitative implementation layer that operationalizes the requirements of existing biorisk management standards. This section maps the present methodology to two key international standards: ISO 35001:2019 (Biorisk management for laboratories) and ISOl/TS 5441:2024 (Competence requirements for biorisk management advisors). Qualitative biorisk standards (ISO 35001) emphasize Plan-Do-Check-Act (PDCA) methodology but lack quantitative metrics like the present log-risk framework.

### 6.1. ISO 35001:2019 – Biorisk Management Systems

ISO 35001:2019 specifies requirements for a biorisk management system, including processes to identify, assess, control, and monitor risks associated with hazardous biological materials [1]. The standard is performance-based, requiring organizations to evaluate control effectiveness and demonstrate continual improvement, but provides limited guidance on quantitative implementation.

The proposed framework directly addresses this gap. The 5-state Markov chain models the entire risk pathway from normal operation to disaster, satisfying the requirement for systematic risk identification. Log-risk scoring (*L* = −log_10_ *P*) provides an objective, comparable metric for risk assessment. Intervention effect functions quantify the impact of training, maintenance, and inspection, enabling evidence-based risk treatment decisions. Bayesian updating with local incident data creates an auditable record of risk evolution, fulfilling monitoring and documented information requirements. The log-risk scale (0-10) facilitates communication to management as required by ISO 35001 clauses on leadership review and communication.

### 6.2. ISO 15190:2020 – Medical Laboratories: Requirements for Safety

ISO 15190:2020 specifies requirements for safe practice in medical laboratories, including risk assessment, role-specific training, safety inspections, and incident reporting with corrective action [3]. The proposed framework provides the quantitative evaluation layer that the standard implicitly requires but does not specify. The Markov chain model operationalises hazard identification by mapping the escalation pathway from procedural deviation to containment loss; the inspection threshold effect formalises the standard’s binary pass/fail inspection logic; and the MCMC updating pipeline converts the incident and near-miss records that ISO 15190:2020 requires laboratories to maintain into continuously refined posterior risk estimates. The ROI analysis addresses the standard’s proportionality principle by providing an auditable, evidence-based basis for intervention prioritisation.

Table 4 maps the framework capabilities to key requirements of ISO 35001:2019 and ISO 15190:2020. The coverage is comprehensive: every major requirement category of both standards – risk identification, assessment, treatment, monitoring, and management communication – is addressed by at least one framework component. Laboratories operating the framework, therefore, generate the quantitative evidence base needed for formal compliance with both standards as a by-product of routine risk management activity.

**Table 4:** Alignment of framework capabilities with laboratory safety and biorisk management standards.

| Functionality | ISO 35001:2019 | ISO 15190:2020 |
| --- | --- | --- |
| 5-state Markov chain | Risk identification (6.1.2) | Hazard identification (5.3) |
| Log-risk scoring | Risk assessment (6.1.2) | Risk assessment (5.3) |
| Intervention effect functions | Risk treatment (6.1.3) | Training & inspection (5.5, 5.6) |
| ROI analysis | Resource allocation (5.1) | Proportionality principle |
| Bayesian updating | Monitoring & review (9.1, 9.3) | Incident reporting (5.8) |
| Near-miss pipeline | Documented information (7.5) | Corrective action (5.9) |
| 0–10 log-risk scale | Management communication (7.4) | — |

### 6.3. Adaptation to Biosecurity scenarios

The Bayesian framework is very well suited for biosecurity because it addresses the rare event problem, that is the challenge of quantifying risk when actual security breaches (disasters) are extremely infrequent or non present in local historical data. In a traditional frequentist approach, the lack of recorded security breach might be misinterpreted as zero risk. In contrast, this framework utilizes the WHO Risk Group classification as a Bayesian prior to establish a baseline of objective risk in terms of the adverse outcome.

The Markov state structure maps naturally to security scenarios – vulnerability (S1), attempted breach (S2), access gained (S3), and loss (S4) – and can evaluate security-specific interventions such as access control, personnel reliability programs, and intrusion detection systems. This adversary-category structure parallels established biosecurity risk assessment tools, such as Sandia National Laboratories’ Biosecurity Risk Assessment Methodology (BioRAM), which formalizes likelihood and consequence scoring for insider, outsider, and colluding adversary scenarios in laboratory settings [22, 21]. These near-miss indicators map onto standard biosecurity adversary categories, such as insider threats (e.g., anomalous badge access by authorized personnel) and unauthorized access attempts (e.g., failed credential use at containment perimeters), consistent with established biosecurity risk assessment methodologies.

This safety-security integration parallel extends beyond biosafety: recent work combining Bayesian networks with escalation-scenario modeling and cost-benefit analysis has been used to jointly evaluate safety and security protection plans for industrial process facilities, including fireproofing and intrusion-detection measures [28], providing an independent, quantitative instance for the safety-security integration proposed here for biosecurity-relevant laboratory scenarios.

## 7. Discussion

### 7.1. Physical Interpretation

The proposed Markov escalation framework is complementary to classical fault tree analysis (FTA). FTA is a deductive, top-down technique best suited to post-incident forensic investigation: given a disaster that has already occurred, it reconstructs the combinations of component, human, and procedural failures that produced it. Its static, combinatorial logic gates are less suited to modeling ongoing systems with scheduled interventions over time. In contrast, the present framework addresses the prospective question that WHO LBM4 anticipates: quantifying and managing risk *before* an incident occurs, using routinely available operational data–training hours, maintenance adherence, inspection scores–rather than requiring a completed incident to analyze. In practice, the two approaches can be used in sequence: FTA to diagnose the causal pathway of a disaster once it occurs, and the present Markov framework to continuously re-calibrate risk and prioritize interventions in the interim between incidents.

The present model can be interpreted through the lens of reliability engineering. Let *R*(*t*) be the reliability of critical equipment over time. Scheduled maintenance resets *R* to near 1.0, while between maintenance occurrences, reliability decays exponentially with time. The effective reliability over a long period depends on both the frequency of resets and the consistency with which they occur. Gaps allow the decay to continue unchecked, producing disproportionately low average reliability.

If maintenance is scheduled at frequency *f* but only adhered to with probability *a*, the mean time between actual maintenance events is 1*/*(*af*), while the mean duration of gaps is (1 − *a*)*/f*. The exponential gap penalty *e*^−*β*(1−*a*)^ approximates the reliability loss during these gaps, with *β* related to the equipment failure rate.

This physical interpretation grounds the model in observable phenomena and provides a basis for estimating *β* from equipment failure data when such are available. Therefore, the model strongly favors data-driven decision making.

It is important to emphasize that the intervention effect magnitudes (e.g., Δ*L*_max_ for training and maintenance, the inspection threshold value, and the exponential gap-penalty coefficient *β*) and the cost parameters used in the ROI analysis (including disaster cost, training cost per hour, and inspection cost) are judgment-based calibrations informed by the epidemiological literature reviewed in Section [2], rather than parameters fitted to facility-specific empirical data. Consequently, the numerical results reported should be interpreted as illustrative demonstrations of the framework’s internal consistency and decision logic, not as validated point predictions for any specific laboratory. Absolute magnitudes will shift under alternative, equally defensible parameter choices; the qualitative patterns (adherence dominating frequency, diminishing training returns, inspection threshold effects) are the framework’s more robust contribution.

### 7.2. Design Choices and Limitations

The proposed framework has some simplifying limitations and design choices:

1. **Discrete state space**: A deliberate design choice was the space of 5 states for the Markov Chain. Since near misses are seldom reported this provides some reasonable idealization of the lab reality.
2. **Stationarity**: The model further assumes transition probabilities that are constant over time. This is dictated by the categorical type of input data (i.e. occurrences of near misses or incidents) and the discrete dynamics.
3. **Independence assumptions**: Interventions are treated as approximately additive in log-space. In reality, interactions may exist (e.g., well-trained staff may compensate for poor maintenance). The current model does not capture such synergies or antagonisms.

Meaningful Bayesian updating requires incident or near-miss data. For ultra-safe laboratories with zero events over many years, posterior distributions remain diffuse.

The current model assumes a single maintenance program affecting all equipment items uniformly. In practice, different equipment items may have different optimal frequencies and different sensitivities to gaps. This was a deliberate design choice in order to simplify presentation and interpretation of results. The present model can be extended by introducing equipment-specific parameters *β*_*i*_ and aggregating effects. This can ultimately enable creation of a complete digital twin of a given lab.

Additionally, the current formulation does not distinguish between preventive and corrective maintenance; this distinction could be incorporated in future versions as well.

The numerical values for maximum effects (Δ*L*_max_ = 2.0 for training, 1.2 for maintenance) are calibrated judgmentally, not empirically derived. While the Bayesian framework can update these, initial priors matter.

The inspection channel is currently modeled as a binary step function: scores below 70 yield no risk reduction, while scores at or above 70 yield the full effect. This formulation mirrors regulatory pass/fail inspection logic, where compliance determinations are typically issued as discrete outcomes rather than continuous scores. However, it does not capture the graduated dynamics of real inspection processes, in which a facility scoring 68 is plausibly closer in underlying safety performance to one scoring 72 than the step function implies. A natural refinement is to replace the hard threshold with a smoothed sigmoid or logistic function centered at the regulatory cutoff, whose steepness could itself be calibrated to reflect how strictly a given regulatory regime enforces the threshold: a very steep curve recovers the current step-function behavior, while a shallower curve yields a gradual transition consistent with continuous inspection quality. We retain the binary formulation in the present work for its transparency and direct correspondence to existing practice, and identify the logistic generalization as a priority for future calibration once paired inspection-score/outcome data become available to estimate the transition steepness empirically.

The additive treatment of intervention effects in log-space assumes conditional independence between training, maintenance, and inspection channels. This choice follows an Occam’s razor principle: in the absence of empirical data quantifying cross-channel interactions (e.g., whether well-trained staff partially compensate for maintenance gaps, or whether intensive training schedules crowd out attention to routine upkeep), introducing interaction terms would add free parameters without a corresponding evidence base to constrain them, risking overfitting to assumptions rather than data. The framework’s stated purpose is to establish auditable baseline risk estimates and decompose intervention contributions in a transparent, reproducible manner, not to model the full joint dynamics of laboratory safety behavior with certainty. Independence is therefore treated as a reasonable default rather than a claim of empirical fact: it yields a conservative, interpretable starting point that a facility can refine as local interaction effects, if any, become evident from accumulated incident and near-miss data. Should future empirical work identify systematic and quantifiable synergies or antagonisms between channels, the additive decomposition can be extended with correction terms informed by that evidence, rather than a priori.

Despite these simplifying assumptions, the model captures the essential non-linearities that govern real maintenance effectiveness and provides a rigorous basis for the adherence-beats-frequency insight that emerges from the analysis.

### 7.3. The data problem

The present paper relies on synthetic BSL-3 scenarios rather than real-world incident logs, a choice that reflects a structural data gap endemic to biosafety as a field rather than a shortcoming of the Bayesian framework itself. The combination of qualitative, categorical risk assessment traditions with the inherent rarity of the events being assessed has historically produced incident records that are sparse and inconsistent across institutions, and that lack the procedural denominators needed for quantitative calibration of any kind, empirical or Bayesian [12, 16]. A recent systematic review by Dhawan et al. analysed 250 reports comprising 712 total cases of laboratory-acquired infection worldwide, finding 276 infections and eight deaths in research laboratory settings and 227 infections and five deaths in clinical laboratory settings, with needle-stick injuries and ineffective use of personal protective equipment or containment measures identified as leading risk factors across both contexts [26]. This case series, like its predecessors [10, 15], still lacks the procedure-level granularity required to estimate per-procedure risk, underscoring a gap in international incident registries that has long been recognised in the literature [15]. It is exactly this combination – rare events, categorical risk assessment, and absent procedural denominators – that makes a Bayesian approach the natural tool of choice, since it updates risk estimates coherently even where frequentist calibration remains structurally out of reach. The results presented here should accordingly be read as a proof-of-concept demonstration of the framework’s internal logic, identifiability properties, and decision-support capabilities, ready for empirical grounding as better data become available. Systematic reviews, such as that of Dhawan et al. [26], and evolving registries, such as that maintained by ABSA International [15], are steadily converging toward denominator-complete datasets, and this trajectory positions the present framework well for direct empirical validation in the near term – reinforcing, rather than undermining, the case for quantitative, Bayesian approaches as the field’s reporting infrastructure continues to mature.

This challenge is not unique to biosafety: even in industries with substantial incident datasets, such as steel manufacturing, Bayesian Network and machine-learning classifiers trained on 5,919 incident records still achieve zero correct classifications for rare high-consequence outcomes such as fatalities, underscoring that data volume alone does not resolve the rare-event estimation problem [27].

### 7.4. Disaster costs

Available evidence indicates that laboratory disaster costs vary widely not only in magnitude but also in composition across jurisdictions. In the United States, cost structures are heavily shaped by litigation and regulatory enforcement: OSHA fines alone have ranged from $31,875 for a fatal academic incident at UCLA to $907,253 across 21 citations for a commercial testing laboratory, while associated litigation and remediation costs have reached $4.5-24.5 million in a single case and $6.7 million in another [23]. By contrast, in China, where 113 documented laboratory accidents resulted in 99 fatalities since 2000, cost accounting has centred primarily on casualty and property-damage statistics rather than litigation or regulatory penalties, reflecting a markedly different liability and enforcement environment [24]. More broadly, cross-national comparisons of work-related accident costs show that methodologies, insurance structures, and reporting conventions for the economic burden of occupational incidents differ substantially between countries, such that a cost estimate calibrated to one regulatory and insurance environment may not transfer directly to another [25]. Laboratories applying the present framework in jurisdictions with different litigation exposure, insurance markets, or regulatory penalty regimes should therefore substitute a locally derived disaster cost estimate rather than relying on the historical U.S.-centric value used here.

### 7.5. Application domains

The proposed framework has distinct but interconnected implications across biosafety and biosecurity. In **biosafety**, it transforms practice from static compliance to dynamic risk management. Log-risk scores enable objective benchmarking between laboratories, allowing a BSL-3 facility operating at *L* = 7.2 to be directly compared to a peer at *L* = 6.8–a difference of 0.4 logs representing a factor of 2.5 in disaster probability. Biosafety officers can calculate marginal ROI for competing interventions, moving from intuition-based to evidence-based resource allocation. The maintenance model’s finding that adherence dominates frequency has immediate practical application: laboratories should focus on executing scheduled maintenance consistently rather than simply increasing frequency. The intuitive 0-10 log-risk scale bridges communication gaps between technical staff and senior management.

For **biosecurity**, the Bayesian framework is well suited to quantifying adversarial rare events where frequentist approaches fail. It should be noted that while the present framework is theoretically compatible with security scenarios, the current quantitative calibrations and intervention functions are primarily optimized for biosafety pathways. Consequently, the formal adaptation, empirical validation, and parameter tuning of the framework specifically for biosecurity applications are subjects for further work. The risk posterior can be further dynamically refined using a near-miss paradigm. For example one could include lower-severity recorded events, such as unauthorized badge swipes, failed attempts to access restricted digital files, or identified vulnerabilities during security audits. This would transform the absence of evidence high severity security incident into a measurable state of uncertainty, honestly reflecting that a lack of incidents is not evidence of the lack of risk. Such an approach would eventually integrate biorisk management as envisioned by ISO 35001, where safety and security investments are optimized within a unified quantitative framework.

## Acknowledgments

This research did not receive any specific grant from funding agencies in the public, commercial, or not-for-profit sectors.

## Disclosure Statement

The authors declare that they have no competing interests.

## Declaration of generative AI and AI-assisted technologies in the manuscript preparation process

During the preparation of this work the author used Claude Sonnet and Deepseek LLMs in order to design proof of concept Python code. The author reviewed and edited the code as needed and takes full responsibility for the computational outcomes.

## Appendix A. Model Specification and Parameter Choices

### Appendix A.1. Log-Space Representation

Let *P* be the probability of a laboratory-acquired infection (disaster) per procedure. The **per-procedure log-risk** *L* is defined as the base-10 logarithm of the conditional mean first-passage time (cMFPT) to state *S*_4_, computed from the forward-escalation sub-chain rooted at state *S*_0_:

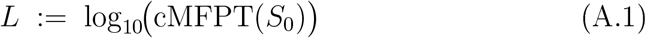

where cMFPT(*S*_*i*_) denotes the mean number of procedures until absorption in *S*_4_ starting from *S*_*i*_, restricted to the sub-chain *{S*_*i*_, *S*_*i*+1_, …, *S*_4_*}* (deescalation probability mass below *S*_*i*_ is removed and rows are renormalised). This restricted sub-chain isolates the forward-escalation channel specific to the intervention class rooted at *S*_*i*_, making *L* sensitive primarily to training while being relatively insensitive to maintenance or inspection.

For reporting purposes the raw one-step disaster probability is also computed:

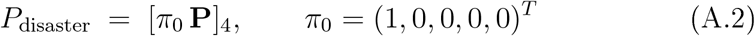

Both quantities increase with improved safety. For the typical operating range, *L* and − log_10_(*P*_disaster_) track each other closely (within ∼0.1 log units).

Two channel-specific companion metrics are defined analogously:

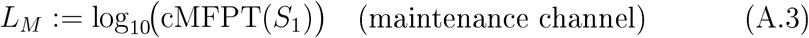

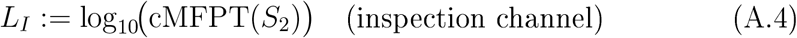

Together (*L, L*_*M*_, *L*_*I*_) form a three-axis decomposition of risk enabling clean, non-overlapping ROI calculations for each intervention type.

### Appendix A.2. WHO Classification as Bayesian Prior

The WHO Risk Group provides a scientifically validated starting point. For a RG-3 agent:

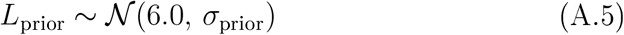

where *σ*_prior_ ≈ 0.15-0.3 logs (factor of 1.4-4 in probability space). This prior is updated as local data accumulate. The mapping RG-1 (*L* ≈ 2), RG-2 (*L* ≈ 4), RG-3 (*L* ≈ 6), RG-4 (*L* ≈ 8) is a modelling choice calibrated to produce disaster probabilities of 10^−2^-10^−8^ per procedure; it is not a WHO-endorsed numerical specification.

### Appendix A.3. Markov Chain State Structure and Edge Classes

Laboratory operations are modelled as a 5-state Markov chain (Fig. 1). The transition matrix **P** (5×5) gives probabilities of moving between states per procedure; row *i* sums to 1 and state *S*_4_ is absorbing.

The implemented edge set covers three distinct failure origin classes:

- **Procedural** (*S*_0_ → *S*_1_, *S*_1_ → *S*_0_, *S*_1_ → *S*_4_, class *H*/*HE*): human-performance primary; the dominant LAI pathway [12].
- **Equipment** (*S*_0_ → *S*_2_, *S*_1_ → *S*_2_, *S*_2_ → *S*_3_, *S*_2_ → *S*_0_, class *E*): maintenance-dominated; includes spontaneous equipment failure *S*_0_ → *S*_2_ (base rate 10^−6.5^, three orders below the procedural path).
- **Containment barrier** (*S*_2_ → *S*_4_, *S*_3_ → *S*_4_, class *EO*): combined equipment and organisational barrier, affected by maintenance and inspection jointly.
- **Uncontrolled de-escalation** (*S*_2_ → *S*_1_, *S*_3_ → *S*_2_, class −): no intervention weight.
- **Natech exogenous bypass** (*S*_0_ → *S*_4_): natura/industrial hazard; parameterised by site profile only, no operational intervention weight.

To improve transparency, each edge carries a cause-class tag linking it to the dominant mechanism it represents:

- *H* (Human): training only;
- *E* (Equipment): maintenance only;
- *HE* (dual Human-Equipment): training and maintenance;
- *EO* (dual Equipment-Organisational): maintenance and inspection.

Inspection acts exclusively on *EO*-tagged edges, reinforcing the final containment barrier on paths closest to *S*_4_.

The log-space update for each transition is:

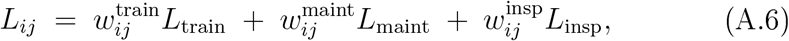

where 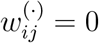 whenever *κ*(*i, j*) is inconsistent with that intervention (strictly sparse weight matrix).

### Appendix A.4. Training Effect Function

Training exhibits diminishing returns (Hill function):

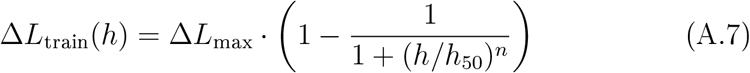

Parameters: Δ*L*_max_ = 2.0 logs (expert training reduces human error up to factor of 100; plausible given empirical 10-100× error reduction between novice and expert [7]); *h*_50_ = 50 h (approximately one week of intensive training); *n* = 2.5 (S-curve matching empirical learning curves [8]). Results are robust to *±*20 % variation in *h*_50_ or *n*.

### Appendix A.5. Maintenance Effect Function

Maintenance effectiveness depends on scheduled frequency *f* (per month) and schedule adherence *a* (proportion completed):

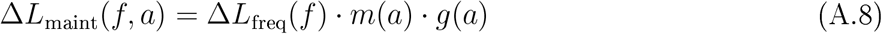

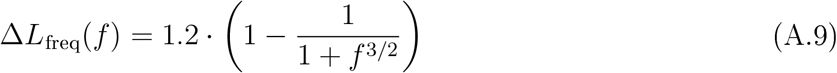

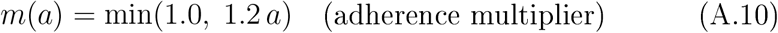

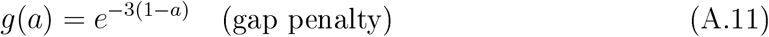

The frequency Hill function saturates at Δ*L*_max_ = 1.2 logs (equipment-related risk reduction of factor ∼16 at optimum; less than training because equipment failures are one of several pathways). Half-saturation at *f* = 1.0/month matches typical BSL-2/3 inspection cycles [19].

The adherence multiplier with synergy factor *α* = 1.2 encodes that consistent partial adherence is slightly more valuable than the raw proportion, reflecting a functional maintenance culture. The min(1.0) cap ensures no super-unit benefit.

The gap penalty *g*(*a*) = exp[− 3(1 − *a*)] reflects Poisson survival *P* (no failure | *k*(1 − *a*)) where *k* = *λt* ≈ 3 expected failures per missed session. This captures accelerating deterioration during gaps:

- *a* = 0.9 (10 % missed): 26 % penalty
- *a* = 0.8 (20 % missed): 45 % penalty
- *a* = 0.7 (30 % missed): 59 % penalty
- *a* = 0.5 (50 % missed): 78 % penalty

The most sensitive parameter is the gap-penalty coefficient *β* = 3; least sensitive is the saturation frequency *f*_0_.

### Appendix A.6. Inspection Effect Function

Inspection produces a binary threshold effect:

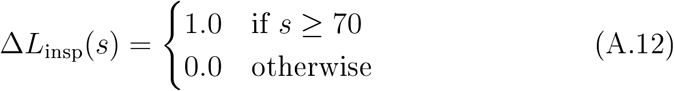

where *s* is the inspection score (0-100). The threshold at 70/100 reflects that inspections either pass or fail a quality standard; incremental improvements below the threshold produce negligible systemic benefit, consistent with the binary pass/fail logic of regulatory biosafety audits.

The 1.0-log input is distributed through the *EO* weight structure (coefficients 0.6 each on *S*_2_ → *S*_4_ and *S*_3_ → *S*_4_), producing a net system-level gain of Δ*L*_*I*_ = 0.600 logs – a factor of ∼4 reduction in conditional disaster probability from the serious or critical states. This allocation explains why the ROI tables report 0.600 rather than 1.0.

### Appendix A.7. Combined Log-Space Update and Risk Calculation

Effects are additive in log-space. The total improvement for transition (*i, j*) is:

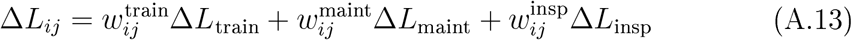

The updated transition probability is:

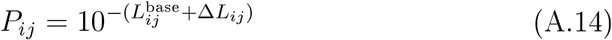

Starting from *π*_0_ = (1, 0, 0, 0, 0)^*T*^, the per-procedure disaster probability is:

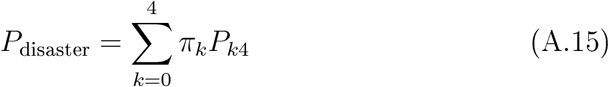

Near-misses are defined as procedures terminating in *S*_1_ – the most observable and reportable safety-relevant events.

### Appendix A.8. Bayesian Likelihood and Prior Distributions

The framework tracks three observable event types over *N* procedures:

- **Near-misses** *n*: rate *p*_01_(*θ*) = [*π*_0_**P**(*θ*)]_1_
- **Light incidents** *ℓ*: rate *p*_23_(*θ*) = *P*_23_(*θ*)
- **Disasters** *d*: rate 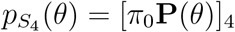

Light incidents substantially improve identifiability of maintenance and inspection parameters compared to a disaster-only likelihood. The joint likelihood is:

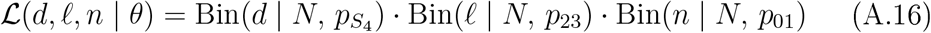

where *θ* = (*h, f, a, s*).

Prior distributions reflect typical operational circumstances (Table A.5):

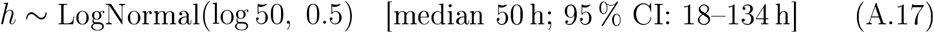

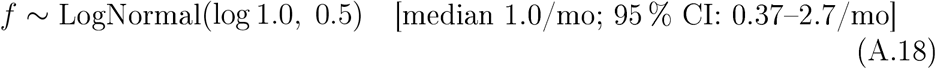

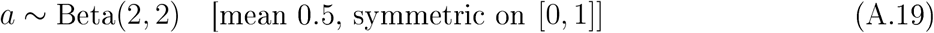

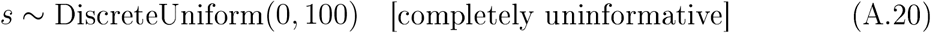

**Table A.5:** Prior distributions and literature basis.

| Parameter | Prior | Literature basis | Source | Notes |
| --- | --- | --- | --- | --- |
| Training | LN(50 h) | 2 h initial +<br>1–2 h annual ×<br>25 y | [17, 18] | Career total |
| Maint. freq. | LN(1.0/mo) | Monthly BSL-2/3 checks | [19] | Standard |
| Maint. adh. | Beta(2,2) | 40–60 % compli-<br>ance | [20] | Conservative |
| Inspection | U(0,100) | Audit scores | – | Ignorance<br>prior |

### Appendix A.9. Cost-Effectiveness Metrics

For an intervention costing *C* that changes log-risk from *L*_0_ to *L*_1_:

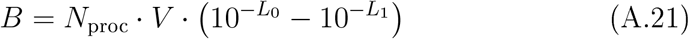

where *N*_proc_ = 10^4^ procedures per year and *V* = $50M (direct – indirect – intangible costs of a laboratory disaster, conservative estimate [9]). Two metrics characterise cost-effectiveness:

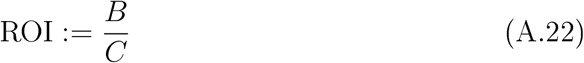

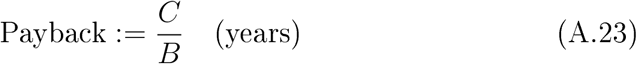

Intervention costs assumed: $100/hour for training (instructor, staff time, materials); $10,000 per 1.0 increment in maintenance frequency (additional staff, parts); $30,000 to cross the inspection threshold (consulting, facility upgrades). These are order-of-magnitude estimates adjustable per laboratory.

### Appendix A.10. Intervention Weight Matrix

Design rationale.

- **Training weights** act on human-performance transitions: procedural deviation onset (*S*_0_ → *S*_1_, *w* = 0.3), procedural recovery (*S*_1_ → *S*_0_, *w* = 0.5), and deviation-to-exposure (*S*_1_ → *S*_4_, *w* = 0.3, dual class *HE*). Training cannot reduce equipment or organisational failure rates.
- **Maintenance weights** target equipment deterioration: spontaneous failure (*S*_0_ → *S*_2_, *w* = 0.8), escalation steps (*S*_1_ → *S*_2_, *S*_2_ → *S*_3_, *w* = 0.8), recovery (*S*_2_ → *S*_0_, *w* = 0.8), and equipment-path exposure (*S*_2_ → *S*_4_, *w* = 0.6; *S*_3_ → *S*_4_, *w* = 0.5; and *S*_1_ → *S*_4_, *w* = 0.5, dual class *HE*).
- **Inspection weights** act exclusively on class *EO* transitions proximate to *S*_4_: *S*_2_ → *S*_4_ (*w* = 0.6) and *S*_3_ → *S*_4_ (*w* = 0.6). Inspection reinforces the final containment barrier by ensuring systemic oversight is operational at the most critical escalation steps. It does not directly affect procedural or equipment failure rates deeper in the chain.
- **Uncontrolled edges** (*S*_2_ → *S*_1_, *S*_3_ → *S*_2_) carry no intervention weight; they represent spontaneous partial recovery not amenable to the three modelled interventions.
- **Natech** (*S*_0_ → *S*_4_) carries no weight; it is exogenous by definition and controlled only by site selection and structural engineering, not operational practice.

## Appendix B. Code Availability

The framework is implemented in Python. The proof-of-concept implementation will be made available upon acceptance as this is a personal repository.

**Table A.6:**
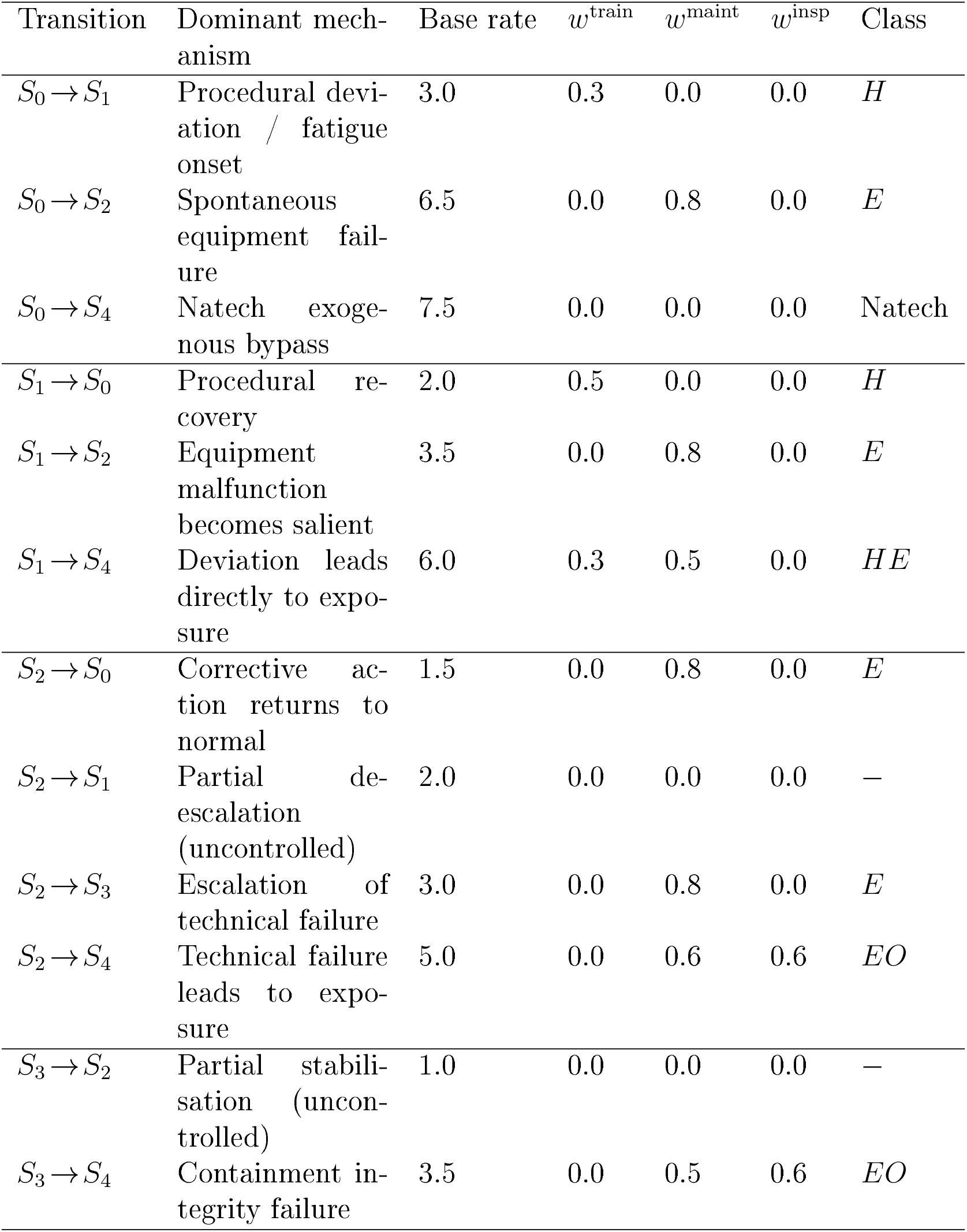
Intervention weight matrix showing cause-class assignments and nonzero weights for the 5-state chain. Base rates are given as − log_10_ 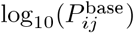.

## Notes

https://github.com/dprodanov/biorisk

### Competing Interest Statement

The authors have declared no competing interest.

### Summary of Updates

Added Wored Example section; Added subsection Adaptation to Biosecurity scenarios; Discussion was broadened - specifically on the part of the data problem and disaster costs; Updated bibliography

## References

[1] International Organiztation for Standardiztation (ISO). ISO 35001:2019_Biorisk management for laboratories and other related organisations. ISO, Geneva, 2019. Available: https://www.iso.org/standard/71293.html.

[2] International Organization for Standardization (ISO). ISO/TS 5441:2024 – Competence requirements for biorisk management advisors. ISO, Geneva, 2024.

[3] International Organization for Standardization (ISO). ISO 15190:2020 – Medical laboratories_Requirements for safety. ISO, Geneva, 2020.

[4] World Health Organization (WHO). Laboratory biosafety manual, 3rd ed. WHO, Geneva, 2004. WHO reference: WHO/CDS/CSR/LYO/2004.11; ISBN 92 4 154650 6. Available: https://www.who.int/publications/i/item/9241546506.

[5] World Health Organization (WHO). Laboratory biosafety manual, 4th ed. WHO, Geneva, 2020. Available (PDF): https://medicine.nus.edu.sg/medphc/wp-content/uploads/sites/33/2023/06/Manual-WHO-Biosafety_4th_edition.pdf.

[6] Reason, J. Human Error. Cambridge University Press, Cambridge, 1990.

[7] Ericsson, K. A., and Charness, N. Expert performance: its structure and acquisition. American Psychologist, 48(8), 1993, 725–747.

[8] Wright, T. P. Factors affecting the cost of airplanes. Journal of the Aeronautical Sciences, 3(4), 1936, 122–128.

[9] Kaplan, S., and Garrick, B. J. On the quantitative definition of risk. Risk Analysis, 1(1), 1981, 11–27.

[10] R.M. Pike, “Laboratory-associated infections: summary and analysis of 3921 cases,” Health Lab. Sci., vol. 13, no. 2, pp. 105–114, 1976.

[11] E.J. Baron and J.M. Miller, “Bacterial and fungal infections among diagnostic laboratory workers: evaluating the risks,” Diagn. Microbiol. Infect. Dis., vol. 60, no. 3, pp. 241–246, 2008.

[12] H. Peng, M. Bilal, and H.M.N. Iqbal, “Improved biosafety and biosecurity measures and/or strategies to tackle laboratory-acquired infections and related risks,” Int. J. Environ. Res. Public Health, vol. 15, no. 12, p. 2697, 2018.

[13] D. Walker and A. Campbell, “A survey of infections in United Kingdom laboratories, 1994-1995,”-J. Clin. Pathol., vol. 52, no. 6, pp. 415–418, 1999.

[14] N.R. Grist and J.A. Emslie, “Infections in British clinical laboratories, 1988-1989,” J. Clin. Pathol., vol. 44, no. 8, pp. 667–669, 1991.

[15] D. Gillum, P. Krishnan, and K. Byers, “A searchable laboratory-acquired infection database,” Appl. Biosaf., vol. 21, no. 4, pp. 203–207, 2016. ABSA International LAI Database: https://my.absa.org/LAI.

[16] R.A. Weinstein and K. Singh, “Laboratory-acquired infections,” Clin. Infect. Dis., vol. 49, no. 1, pp. 142–147, 2009. DOI: 10.1086/599104.

[17] University of Nevada, Reno, Chapter 6: Laboratory Training – Biosafety Manual, 2023.

[18] Boston University, Chapter 6: Laboratory Training – Office of Research, 2016.

[19] Qualia Bio, BSL-3 Lab Maintenance Schedule Template 2025, 2025.

[20] M. A. Faber et al., “Best practices for implementing biosafety inspections in a decentralized institutional biosafety committee,” PLOS ONE, vol. 18, no. 10, e0292940, Oct. 2023.

[21] S. Caskey and P. Edgar, Biosecurity Risk Assessment Methodology (BioRAM) v. 2.0, [Computer Software], Dec. 2019. Available: 10.11578/dc.20220414.20.

[22] S. Caskey, J. Gaudioso, Shigematsu, G. Risi, E. Prat, R. Salerno, S. Wagener, J. Kozlovac, and V. Halkjaer-Knudsen, Biosafety Risk Assessment Methodology, Sandia National Laboratories, SAND2010-6487, October 2010

[23] R&D World, “The price of a lab accident: When safety failures can run in the millions,” R&D World, 2026 Available: https://www.rdworldonline.com/the-price-of-a-lab-accident-when-safety-failures-can-run-in-the-millions/

[24] Y. He, N. S. Kuai, L. M. Deng, Z. L. Wang, and M. J. Peng, “An investigation into accidents in laboratories in universities in China caused by human error: A study based on improved CREAM and SPAR-H,” Heliyon, vol. 10, no. 7, p. e28897, 2024.

[25] D. Elsler, J. Takala, and J. Remes, “An international comparison of the cost of work-related accidents and ill health,” European Agency for Safety and Health at Work (EU-OSHA) report, 2017.

[26] S. Dhawan, A. G. Muluneh, W. Pan-Gnum, C. R. MacIntyre, K. Summermatter, and S. D. Blacksell, “Risk factors and mitigation strategies of laboratory-acquired infections in research and clinical laboratories worldwide: a systematic review,” Lancet Microbe, vol. 6, no. 9, p. 101157, 2025.

[27] S. Pandey, A.K. Singh, S. Parhi, S. Jha, Towards safer steel operations with a multi-model framework for improved accident prediction, Scientific Reports 15 (2025) 13293. 10.1038/s41598-025-96028-0

[28] G. Marroni, S. Kuipers, G. Landucci, V. Casson Moreno, Probabilistic Evaluation of Integrated Safety-Security Protection Plans for Process Facilities, Risk Analysis 46 (4) (2026) e70230. 10.1111/risa.70230

[29] L. C. Klotz, E. J. Sylvester, The unacceptable risks of a man made pandemic, Bulletin of the Atomic Scientists (2012). http://thebulletin.org/unacceptable-risks-man-made-pandemic (Accessed June 10, 2023).

